# Algorithm for the Quantitation of Variants of Concern for Rationally Designed Vaccines Based on the Isolation of SARS-CoV-2 Hawaiʻi Lineage B.1.243

**DOI:** 10.1101/2021.08.18.455536

**Authors:** David P. Maison, Lauren L. Ching, Sean B. Cleveland, Alanna C. Tseng, Eileen Nakano, Cecilia M. Shikuma, Vivek R. Nerurkar

## Abstract

SARS-CoV-2 worldwide emergence and evolution has resulted in variants containing mutations resulting in immune evasive epitopes that decrease vaccine efficacy. We acquired clinical samples, analyzed SARS-CoV-2 genomes, used the most worldwide emerged spike mutations from Variants of Concern/Interest, and developed an algorithm for monitoring the SARS-CoV-2 vaccine platform. The algorithm partitions logarithmic-transformed prevalence data monthly and Pearson’s correlation determines exponential emergence. The SARS-CoV-2 genome evaluation indicated 49 mutations. Nine of the ten most worldwide prevalent (>70%) spike protein changes have *r-*values >0.9. The tenth, D614G, has a prevalence >99% and *r*-value of 0.67. The resulting algorithm is based on the patterns these ten substitutions elucidated. The strong positive correlation of the emerged spike protein changes and algorithmic predictive value can be harnessed in designing vaccines with relevant immunogenic epitopes. SARS-CoV-2 is predicted to remain endemic and continues to evolve, so must SARS-CoV-2 monitoring and next-generation vaccine design.

## Introduction

Since the origin of the Coronavirus Disease 2019 (COVID-19) pandemic, severe acute respiratory syndrome coronavirus-2 (SARS-CoV-2) has rapidly evolved into seven Variants of Interest (VOI) and four Variants of Concern (VOC).^1^ Further, as of July 15, 2021, over 2,559,000 SARS-CoV-2 genomic sequences have been deposited in the publicly available GenBank and the Global Initiative on Sharing Avian Influenza Data (GISAID) databases.^2^ From the establishment of the now universal D614G substitution^3^ to the emergence of the VOC and VOI with dozens of different mutations across their respective genomes,^1^ the SARS-CoV-2 evolution, and adaptations, are apparent and constant. To give nomenclature to these evolutionary events, the Centers for Disease Control and Prevention (CDC) has classified certain lineages as VOC and VOI to denote highly adapted and immunologically evasive strains of SARS-CoV-2.^1^ More recently, the World Health Organization (WHO) has further given its own classification to emerging SARS-CoV-2 lineages using letters of the Greek alphabets.^4^

Fortunately, early in the pandemic, governments and private sectors around the globe poured resources into producing efficacious vaccines. In the United States, three of these vaccines are authorized and recommended by the U.S. Food and Drug Administration (FDA).^5, 6^ Unfortunately, all these vaccines have reduced efficacy against all VOC,^1^ an effect likely to amplify further as the virus evolves significantly from the vaccine design of the original strain. The loss of efficacy can be attributed to the alteration of immunogenic epitopes.^7^ Several of these mutations are found in the spike protein, the protein used in the vaccine design, and therefore allows the virus to evade antibodies targeted to the original strain that vaccines utilize. Similar to the annual influenza virus vaccine, the evolution of SARS-CoV-2 presents the dilemma of how to redesign next-generation vaccines to keep up with the evolution of the virus.

One attempt to match the vaccine to the evolution of the virus was by Moderna. In response to the B.1.351 VOC considerably reducing the efficacy of the Novavax vaccine,^8^ Moderna explored the use of the B.1.351 VOC sequence in their mRNA vaccine design.^9^ Promisingly, the newly adapted vaccine increased neutralization against the B.1.351 VOC when given as a booster.^10^ However, the B.1.351 was only 1.15% prevalent worldwide in April 2021. Therefore, the continuous clinical trial evaluation against emerging VOC is not practical to match the rate and diversity with which mutations and VOC emerge worldwide.

Hawai’i has been disproportionately affected by COVID-19 in terms of race, wherein 20% of the cases occur in 4% of the population of Pacific Islanders.^11, 12^ Understanding the SARS-CoV-2 lineage discrepancy in Hawai’i will allow for a greater understanding of the pandemic’s nature worldwide. Additionally, adapting vaccines to match the nature of the viral sequence may alleviate these discrepancies. All four of the VOC recognized by the CDC are present in Hawai’i.^13^

To answer the dilemma of redesigning next-generation SARS-CoV-2 vaccines, we present and further validate our archetype quantitative analysis^14^ for determining the emergence of individual mutations and variants, alike, as an algorithm. This algorithm is a platform for monitoring the virus and determining appropriate vaccine design as we proceed into the evolution and endemicity of SARS-CoV-2.^15^ We utilized SARS-CoV-2 isolated in Hawai’i, and whole genome sequences (WGS) deposited in GenBank and GISAID, in combination with VOC and VOI to validate the algorithm, a prototype alpha test for rationally-designing logical next-generation vaccines. As SARS-CoV-2 evolves, so must SARS-CoV-2 monitoring and vaccine design.

## Methods

### Patient Samples

Human clinical samples analyzed in this report were part of the University of Hawai’i at Manoa (UHM) approved IRB - H051 study (# 2020-00367) (#NCT04360551). The samples were from two patients^14^ (patient identification [PID] 498 and PID 708) collected as oropharyngeal (OPS) and nasal (NS) swabs at days 5 and 3, following symptom onset, respectively. SARS-CoV-2 positive status was confirmed using quantitative reverse-transcriptase-polymerase chain reaction (qRT-PCR). The samples were stored at -80°C as part of the UHM IBC approved study (# 20-04-830-05).

### Virus Isolation

Virus isolation was conducted using Vero E6 (ATCC CRL-1686) cells and from PID 498 OPS collected in the viral transport medium (VTM), as described previously.^16^ Briefly, following 1 hour infection, cells were monitored for cytopathic effect (CPE) using a Cytosmart Microscope monitoring the same location in the flask. After observing significant CPE at 48 hours, supernatant was blind passaged three times in the Vero E6 cells. Virus isolation was confirmed with plaque assay using a double overlay, performed as previously described.^17^

### Growth Kinetics

Following isolation of SARS-CoV-2, Isolate USA-HI498 2020, a growth kinetics study was conducted by seeding monolayers of Vero E6 cells in 6 well plates at a cell density of 3×10^5^/well one day prior to the assay. For the assay, cells were infected with SARS-CoV-2 USA-WA1/2020 and HI498 2020 isolates. Briefly, on the day of the assay, DMEM in 10% FBS was removed from wells with Vero cells, wells were washed twice with serum-free DMEM, and inoculated with multiplicity of infection (MOI) 0.1 and 1 virus isolates diluted in 500 µL DMEM in 2% FBS and incubated at 37°C and 5% CO_2_ for two hours. Following the two-hour adsorption, the infectious supernatant with virus was removed, and monolayers were washed twice with DMEM in 2% FBS, and further incubated in 2 mL DMEM with 2% FBS at 37°C and 5% CO_2_ until supernatant collection at 0, 12, 24, and 48 hours.^16^

### RNA Extraction, qRT-PCR, and Genomic Equivalence

For determining Genomic Equivalence, RNA extraction was conducted using the QIAamp® Viral RNA Mini Kit (Qiagen, Cat# 52906) following the manufacturer’s instructions and as described previously.^14^ RNA extraction was conducted on eight, ten-fold serial dilutions (10^0^ to 10^-8^) independently.^17^ The primers and probes (N1 set, N2 set, and RdRp set) used are described previously.^18, 19^ A TaqMan® multiplexed qRT-PCR method was used for the N1 and N2 primer sets. The QuantaBio qScript® XLT One-Step qRT-PCR Tough Mix (Cat# 95132) was used to conduct the qRT-PCR on a ABI StepOnePlus™ Real-Time PCR system. A SYBR Green qRT-PCR method was used for the RdRp primer set. The QuantaBio qScript® cDNA Synthesis Kit (Cat# 95047) and QuantaBio PerfeCTa® qPCR ToughMix™, Low ROX™ (Cat# 95114) was used to perform the qRT-PCR on a ABI StepOnePlus™ Real-Time PCR system.

Standard curves for the SARS-CoV-2 isolates by N1, N2, and RdRp genes were produced by plotting cycle threshold (Ct) values against corresponding plaque forming units (PFU) per mL evaluated by plaque assay for the eight ten-fold serial dilutions of SARS-CoV-2 virus isolates.^17^ The standard curve produced from the ten fold-serial dilutions of the virus was used to interpolate the results of the growth kinetics experiment. All qRT-PCR and Genomic Equivalence data were analyzed and visualized using GraphPad Prism 9 Version 9.2.0.

### Whole Genome Sequencing

For WGS, RNA extraction was conducted using the third blind passage with the QIAamp® Viral RNA Mini Kit (Qiagen, Cat# 52906) following the manufacturer’s instructions, as described previously.^14^ Briefly, viral RNA was eluted in 60 µL of the elution buffer. RNA extraction was confirmed using the Takara RNA LA PCR Kit (CAT #RR012A) and previously reported primer sets.^14, 20^ RNA was reverse transcribed into cDNA using the Takara RNA LA PCR Kit (Cat #RR012A) according to the manufacturer’s protocol but with an extension time of 90 minutes. WGS was conducted by the ASGPB Core, UHM. Briefly, libraries prepared as per the manufacturer’s protocol (Illumina Document #1000000025416 v09) using Illumina DNA Prep kit (Cat #20018704) and Nextera XT indexes were sequenced using the MiSeq Reagent Kit v3 (600 cycle) (Cat #MS-102-3003) and an Illumina MiSeq sequencer.

### Informatics

WGS reads were compiled using the UHM MANA High-Performance Computing Cluster (HPC). Raw fastq sequence files were evaluated by the FASTQC program^21^ for technical sequence error. After confirming a lack of technical errors, low-quality sequences were filtered and trimmed from each read with Trimmomatic^22^ using paired-end adapter sequence NexteraPE-PE (ILLUMINACLIP:NexteraPE-PE:2:30:10) and a sliding 4 base window evaluating for quality with a PHRED score over 30. Trimmed result quality was confirmed with FASTQC. The trimmed-paired-end reads were then mapped to the NC_045512 reference genome using Bowtie2^23^ and variants called with samtools mpileup^24^ and transformed from VCF to FASTQ using bcftools and vcfutils^25^ and finally converted to FASTA using seqtk. For comparison and validation, the fastq file was also inputted into Geneious Prime 2021.1.1 to produce FASTA files with the coronavirus assembly workflow.^26^ The resultant consensus sequence was defined as Hawai’i Isolates and submitted to GenBank (SARS-CoV-2, Isolate USA-HI498 2020 (MZ664037) and SARS-CoV-2, Isolate USA-HI708 2020 (MZ664038)). The lineage of each sequence was determined with the Phylogenetic Assignment of Named Global Outbreak (PANGO) Lineage nomenclature.^27–29^

### Hawai’i Sequences and Lineage Searches

All Hawai’i sequences as of July 28, 2021, from both GenBank and GISAID were downloaded and searched for the presence of potential lineages of concern using PANGO lineage as described previously.^13, 27–29^

### Variant Comparison

A comparison was conducted to evaluate the mutations in B.1.243 compared to 12 other VOC, VOI, and variants as of May 12, 2021. The NCBI SARS-CoV-2 resources genomic reference sequence from Wuhan was used to define the S gene (NC_045512).^30^ Each of the sequences underwent pairwise alignment with NC_045512 to define S gene mutations. The Hawai’i Lineage (B.1.243) sequence selections were SARS-CoV-2 HI498 and HI708. Sequences for VOC, VOI, and other variants that garnered attention throughout this pandemic were selected with criteria of earliest complete collection dates with unambiguous S gene sequences. Lineages used were: B.1.1.7 (United Kingdom VOC, EPI_ISL_601443)^31, 32^, B.1.1 (Nigeria variant, EPI_ISL_729975)^33, 34^, B.1.351 (South Africa VOC, EPI_ISL_712081)^35^, B.1.1.298 (Denmark variant, EPI_ISL_616802)^36^, B.1.427 (California VOI, EPI_ISL_1531901)^37^, B.1.429 (California VOI, EPI_ISL_942929)^38^, P.1 (Brazil/Japan VOC, EPI_ISL_792680)^39, 40^, P.2 (Brazil VOI, EPI_ISL_918536)^41^, B.1.617.1 (India VOI, EPI_ISL_1372093)^42^, B.1.617.2 (India VOC, EPI_ISL_1663516)^43^, and B.1.525 (United Kingdom/Nigeria VOI, EPI_ISL_1739895).^44^

### Quantitation of Variants and Amino Acid Substitution/Deletions in Comparison to Epitope Mapping of the Spike Protein

From the aforementioned variant comparison section, the selected S gene sequences underwent pairwise alignment with NC_045512 in SnapGene, and SNPs were identified. The SNPs were inputted into the SnapGene sequence feature and Nextclade^45^ to determine amino acid substitutions (AAS). Non-synonymous substitutions were confirmed in GISAID using the metadata for each accession number.

The PANGO Server was used to identify and confirm the lineage of each of the aforementioned thirteen strains described in the Variant Comparison section.^28, 29^ The lineages and their collective AAS were identified and individually searched within GISAID for worldwide prevalence from March 2020 - April 2021. Each lineage was filtered separately, as were AAS. Parameters for selection were sequences that included a full month, day, and year of collection. Each month’s prevalence for each lineage and AAS was logarithmically transformed and evaluated against month as an interval value with Pearson’s correlation to determine an exponential increase in worldwide emergence as described previously.^14^ Pearson’s was calculated using RStudio version 1.3.1093 (R version 4.0.3) and plotted with the ggplot2 package. The Pearson’s correlations for AAS and lineages were then compared in a corresponding pairwise heat map to evaluate if AAS emergence occurs independently of, or in tandem to, lineage emergence.

Separately, the following search parameter was used in PubMed to locate *in silico* studies predicting vaccine epitopes to SARS-CoV-2: “((B-cell) OR (B cell)) AND ((T-cell) OR (T cell)) AND (peptide) AND (vaccine epitope) AND ((SARS-CoV-2) OR (COVID-19)).” From this search on January 28, 2021, the three most recent articles^46–48^ and the three best matching articles^49–51^ were selected for further analysis by mapping to the Spike protein. All predicted epitopes able to be searched and defined with SnapGene’s “Find Protein Sequence” feature were included. Article overlaps in the systematic review were only included once.

### Algorithm

The algorithm herein described was developed from the quantitation of the ten most emerged amino acid substitutions and deletions. The algorithm is as follows:

if Pearson’s *r*-value ≥ 0.9:

if Previous Month’s Prevalence > 0.3:

Emerging (Mutation of Concern) (Include in Next Generation Vaccine Design)
else if 0.02 ≤ Previous Month’s Prevalence ≤ 0.3:

Emerging (Mutation of Interest)
else:

Not Emerging

else:

if Previous Month’s Prevalence ≥ 0.9:

Emerged (Mutation of Concern) (Include in Next Generation Vaccine Design)
else if Previous Month’s Prevalence > 0.5:

Emerged (Mutation of Interest)
else:

Not Emerged/Emerging

This algorithm classifies AAS and deletions into two categories based on Pearson’s *r-*value, *r*≥0.9 and *r*<0.9. For AAS and deletions to be called out as concerning, the *r*-value should be ≥0.9 and the previous month’s worldwide prevalence of these AAS and deletions should be >30%. Further, these concerning AAS and deletions can be considered for inclusion in the next-generation vaccine design. If the *r*≥0.9 and the previous month’s prevalence is between 2% and 30%, then the AAS or deletion is classified as interesting, and needs to be evaluated in a research setting. The same algorithm can also be applied for standardizing classifications of SARS-CoV-2 lineages as of interest or concerning.

If the *r*<0.9, then the focus is on previous month’s prevalence of AAS and deletions, as after a mutation is established, there is no longer need to evaluate emergence. If the previous month’s prevalence is ≥90%, then the mutation is established in the SARS-CoV-2 genome and should be considered as concerning and be part of the next-generation vaccine. If the previous month’s prevalence is >50%, then the mutation represents the majority, and needs to be considered as interesting and evaluated in a research setting. Again, the same algorithm can also be applied for standardizing classifications of SARS-CoV-2 lineages as of interest or concerning.

### B.1.243 Phylogeny and Origin Tracking

Origin tracking was accomplished as described previously.^13^ The sequences from Hawai’i were obtained through both GenBank and GISAID, along with SARS-CoV-2, Isolate USA-HI498 2020 and SARS-CoV-2, Isolate USA-HI708 2020 (from this study), to accomplish the origin determination.

## Results

### Virus Isolation, Growth Kinetics, and Genomic Equivalence

Virus was isolated from an OPS collected five days following symptom onset from an individual with PCR confirmed SARS-CoV-2 infection and propagated in Vero E6 cells. A stock of the SARS-CoV-2, Isolate USA-HI498 2020 was produced following three blind passages in Vero E6 cells and titered to 1.28 x 10^7^ PFU/mL. Minimal CPE was observed at 12 hours, moderate CPE at 24 hours, and significant CPE at 48 hours (Figure 1 A-D). Plaque assay confirmed SARS-CoV-2, Isolate USA-HI498 virus isolation at 1.28 x 10^7^ PFU/mL and SARS-CoV-2 USA-WA1/2020 at 3.88 x 10^7^ PFU/mL. Viral copy number analysis using N1, N2 and RdRp primers as well as microscopic observation showed no significant differences between the SARS-CoV-2, Isolate USA-HI498 2020 and SARS-CoV-2 USA-WA1/2020 (Figure 1 E-M).

**Figure 1.**
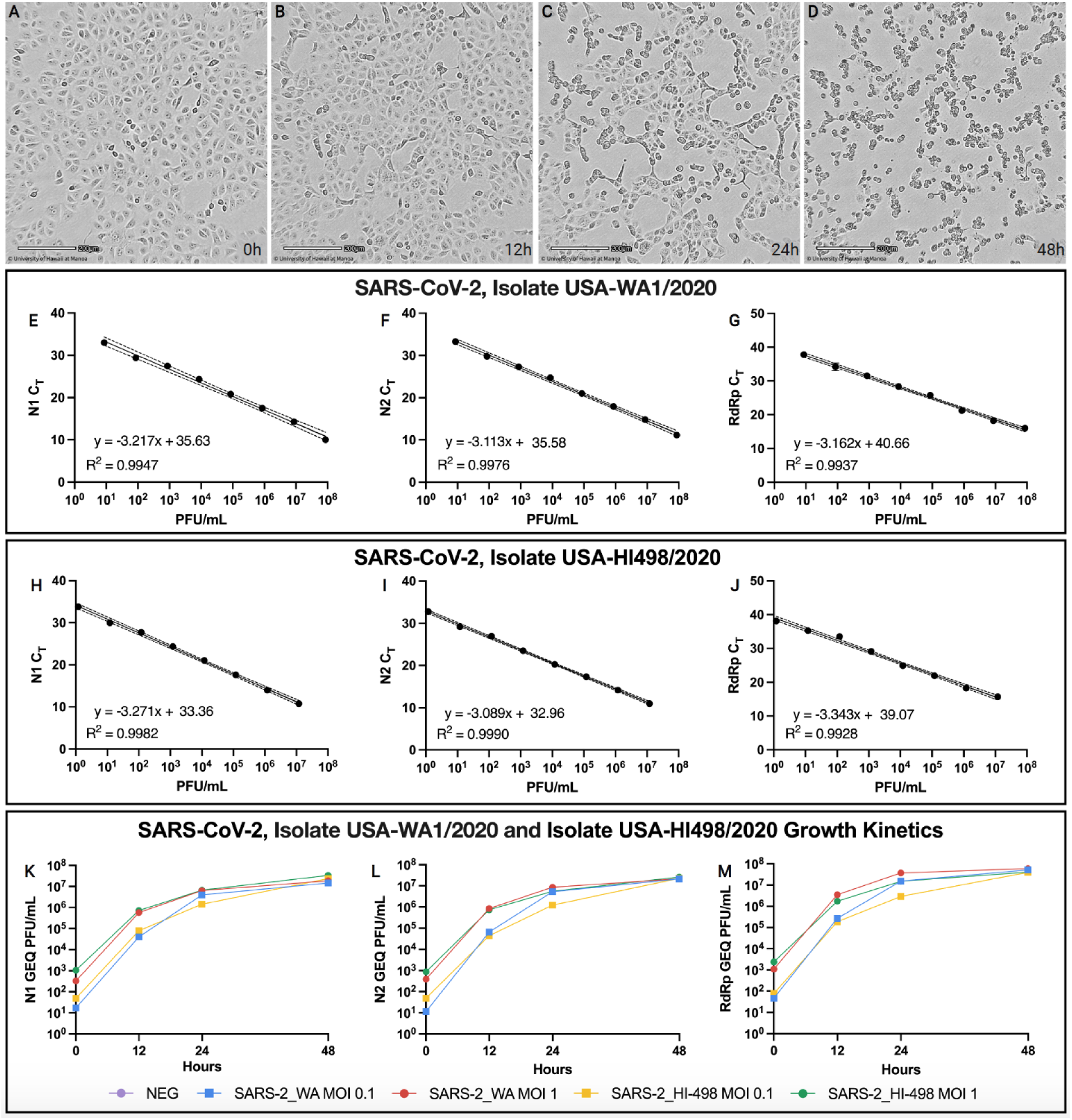
Cytopathic Effect and Growth Kinetics of SARS-CoV-2, Isolate USA-HI498 2020. The figure shows time-lapse cell images of VeroE6 cells infected with SARS-CoV-2 isolate USA-HI498 2020 at different time points, demonstrating the cytopathic effect (CPE) induced by the virus at multiplicity of infection (MOI) 1. A) 0 hr., B) 12 hr., C) 24 hr., and D) 48 hr. Scale bar equals 200 µm. E) Genomic equivalent (GEQ) comparison between SARS-CoV-2 USA-HI498 2020 isolate at MOI 0.1 (yellow) and 1 (green), with the SARS-CoV-2 Washington (WA) isolate at MOI 0.1 (blue) and 1 (red), using N1 (E,H,K), N2 (F,I,L) and RdRp (G,J,M) primers.

### Whole Genome Sequencing and Informatics

RNA extraction was confirmed using RT-PCR as described previously.^14^ There were 702,978 and 792,952 reads for SARS-CoV-2, Isolate USA-HI498 2020 and HI708, respectively (Table 1). FastQC confirmed the quality of both the untrimmed and trimmed fastqc files.

**Table 1.** Genetic Characteristics of the Hawaii SARS-CoV-2 Variant B.1.243 Isolates, USA-HI498 2020* and USA-HI708 2020**

| Gene | Nucleotide |  |  | Amino Acid |  |  |
| --- | --- | --- | --- | --- | --- | --- |
|  | Loci | Wild Type | Mutant | Loci | Wild Type | Mutant |
| 5' UTR | 241 | C | T | - | - | - |
| ORF1ab | 3,037 | C | T | 924 | Phe (F) | Phe (F) |
| ORF1ab | 10,741 | C | T | 3492 | Asp (D) | Asp (D) |
| ORF1ab | 12,076*** | C | T | 3937 | Asn (N) | Asn (N) |
| ORF1ab | 14,408 | C | T | 4715 | Pro (P) | Leu (L) |
| ORF1ab | 20,268 | A | G | 6668 | Leu (L) | Leu (L) |
| S | 23,403 | A | G | 614 | Asp (D) | Gly (G) |
| S | 23,604 | C | A | 681 | Pro (P) | His (H) |
| S | 24,076 | T | C | 838 | Gly (G) | Gly (G) |
| N | 28,854 | C | T | 194 | Ser (S) | Leu (L) |
| N | 29,266*** | G | A | 331 | Leu (L) | Leu (L) |
| 3' UTR | 29,710 | T | C | - | - | - |

### Hawai’i SARS-Cov-2 Sequences and Lineage Searches

From GenBank, 317 full-genome SARS-CoV-2 sequences were obtained on July, 28, 2021. Further, an additional 2,942 sequences were obtained from GSAID. Hawai’i has 52 unique lineages in the 3,259 representative sequences (A.1, A.2.2, A.3, AY.1, AY.2, B, B.1, B.1.1, B.1.1.207, B.1.1.222, B.1.1.304, B.1.1.316, B.1.1.380, B.1.1.416, B.1.1.519, B.1.1.7, B.1.108, B.1.139, B.1.160, B.1.2, B.1.234, B.1.241, B.1.243, B.1.265, B.1.298, B.1.340, B.1.351, B.1.357, B.1.36.8, B.1.369, B.1.37, B.1.400, B.1.413, B.1.427, B.1.429, B.1.517, B.1.526, B.1.561, B.1.568, B.1.575, B.1.588, B.1.595, B.1.596, B.1.601, B.1.609, B.1.617.2, B.1.623, B.6, P.1, P.1.1, P.2, R.1). As of April 12, 2021, GISAID reported 8,809 sequences of B.1.243 lineage worldwide. The B.1.243 variant was represented by 717 of the 1,002 (72%) sequences curated from Hawai’i. This prevalence decreased to 23% by July 28, 2021. Worldwide, GISAID reported 8,809 sequences of B.1.243 lineage.

### Quantitation of SARS-Cov-2 Variants and Amino Acid Substitution/Deletions in Comparison to Epitope Mapping of the Spike Protein

The output S gene alignment between the thirteen genomic sequences identified 49 SNPs. Pearson’s correlation on logarithmically-transformed prevalence was calculated for the twelve SARS-CoV-2 variants in this study in order of highest to lowest *r* value as outlined in Table 2 and Figure 2.

**Figure 2.**
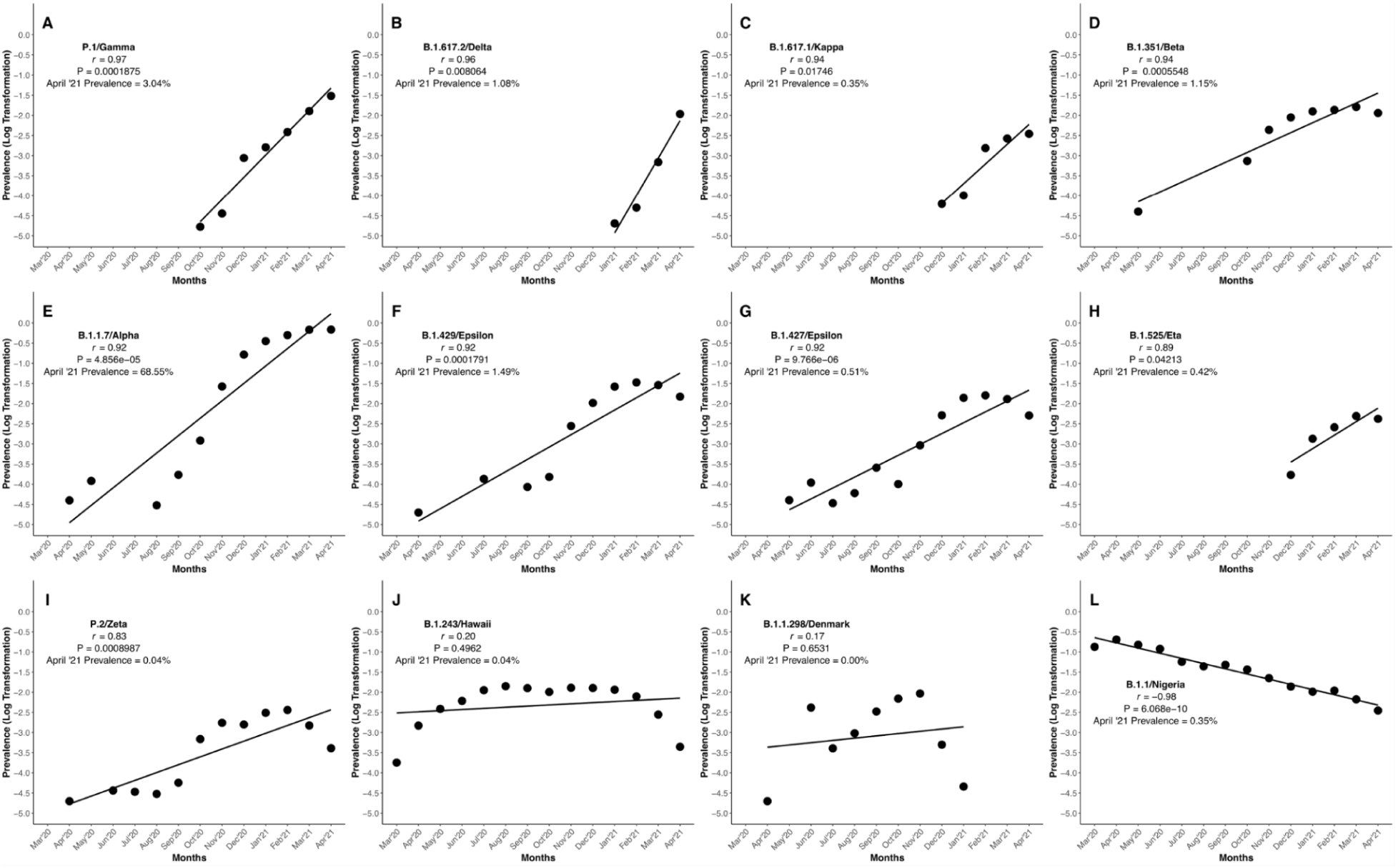
Pearson’s Correlation on Logarithmically-Transformed Prevalence Ratios of SARS-CoV-2 Variant of Concern, Variants of Interest, and other Lineages. This figure demonstrates the quantitation of SARS-CoV-2 variants of concern, variants of interest, and other lineages. The emergence and disappearance of variants/lineages of SARS-CoV-2 is evaluated by Pearson’s correlation of logarithmic transformation prevalence data. Variants are displayed in order of decreasing *r* value (A) P.1, B) B.1.617.2, C) B.1.617.1, D) B.1.351, E) B.1.1.7, F) B.1.429, G) B.1.427, H) B.1.525, I) P.2, J) B.1.243, K) B.1.1.298, and L) B.1.1).

**Table 2.**
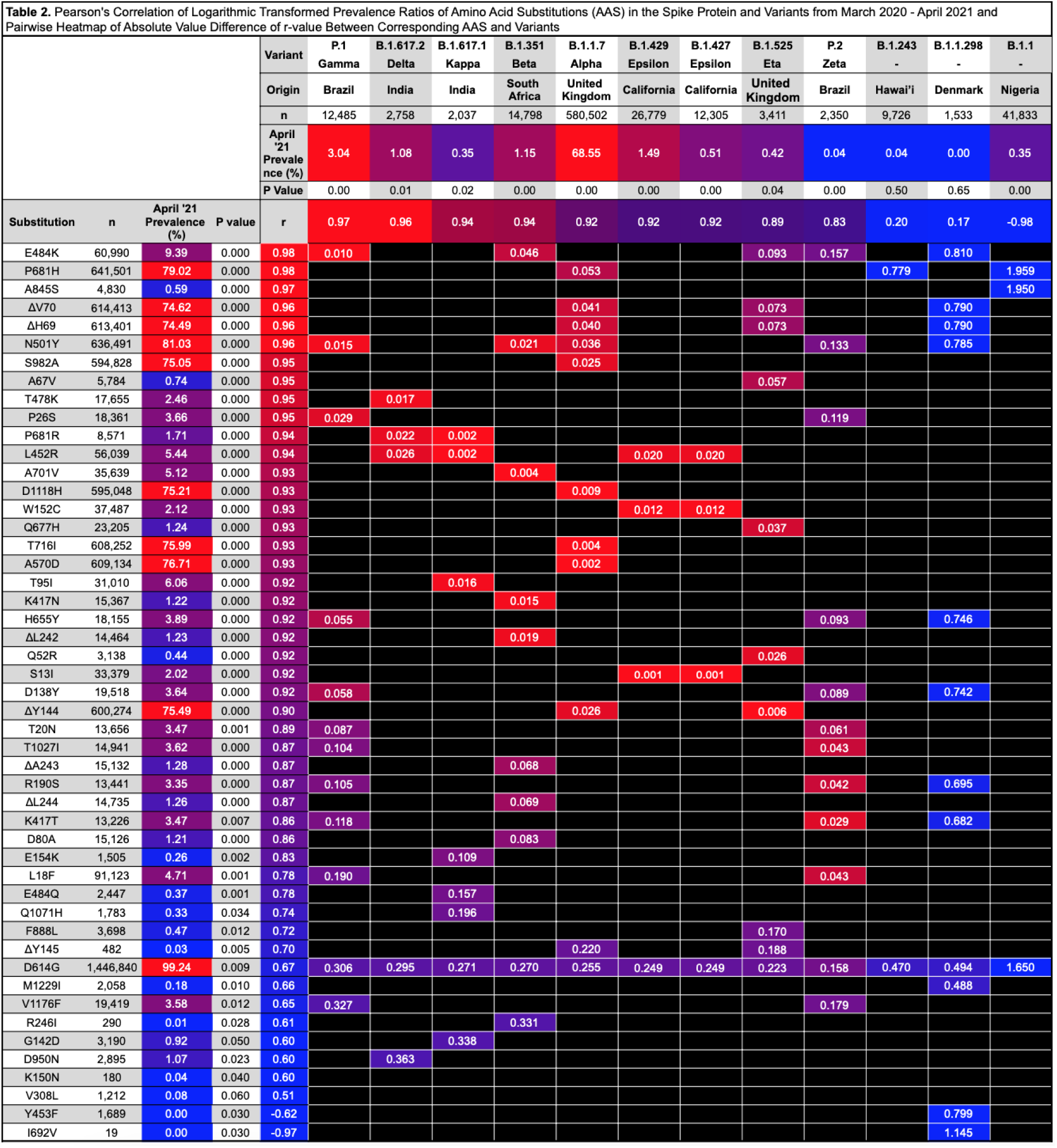
Pearson’s Correlation of Logarithmic Transformed Prevalence Ratios of Amino Acid Substitutions (AAS) in the Spike Protein and Variants from March 2020 - April 2021 and Pairwise Heatmap of Absolute Value Difference of *r*-value between Corresponding AAS and Variants

Further, of the 49 identified SNPs in the S gene, 44 resulted in non-synonymous AAS and deletions in the protein, and 5 were synonymous as outlined in Figure 3A, 3B, 3Ci-xii and Tables 2 and 3. Pearson’s correlation on logarithmically-transformed prevalence was calculated for all identified SARS-CoV-2 AAS and deletions (Tables 2, 3, Figure 4, and Supplementary Figure 1).

**Figure 3.**
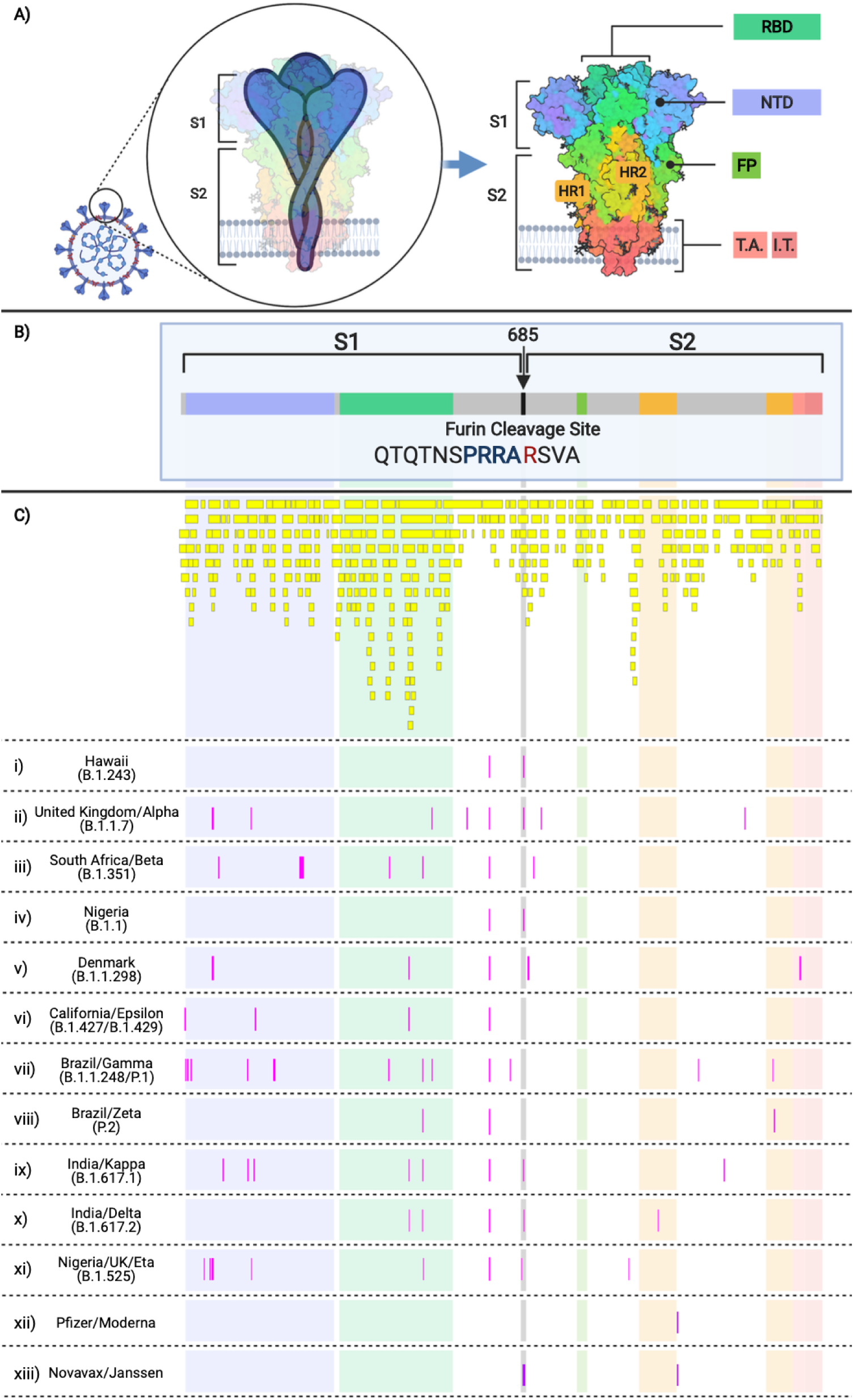
SARS-CoV-2 Spike Protein Domains and Relation to B and T cell Epitopes, Variant Amino Acid Substitutions, and Vaccine Amino Acid Substitutions. This figure demonstrates the evolution of the SARS-CoV-2 variants by depicting the location of the variants substitutions and deletions in the context of spike domains and epitopes. A) Cartoon rendering of SARS-CoV-2 and the 1,273 amino acid long spike protein overlay onto the color-coded crystallographic structure determined by electron microscopy (PBD ID: 6VXX-PDB). The individual protein domains are color-coded: N-terminal domain (NTD) (light purple) (residues 14-305), receptor-binding domain (RBD) (teal green) (residues 319-541), furin (F) (residues 682-685), fusion protein (FP) (green) (residues 788-806), heptad repeat 1 (HR1) (orange) (residues 912-984), heptad repeat 2 (HR2) (orange) (residues 1163-1213), transmembrane anchor (TM) (light pink) (1213-1237), and intracellular tail domain (IT) (dark pink) (1237-1273). B) Two-dimensional layout of the spike protein and domains with the addition of the S1/S2 furin cleavage site (RRA/R) (682-685) (black). C) *In silico* predicted B and T cell epitope loci revealing 393 *in silico* B and T cell epitopes mapped here individually as a yellow boxes i)-xiii) Amino acid substitutions present in the corresponding variant shown in pink boxes in comparison to the reference sequence NC_045512. i) B.1.243 Hawaii; ii) B.1.1.7 United Kingdom; iii) B.1.351 South Africa; iv) B.1.1 Nigeria; v) B.1.1.298 Denmark; vi) B.1.427 and B.1.429 California; vii) P.1 Brazil; viii) P.2 Brazil; ix) B.1.617.1 India; x) B.1.617.2 India; xi) B.1.525 United Kingdom/Nigeria; xii) Pfizer and Moderna mRNA sequences with artificially added substitutions K986P and V987P; xiii) Novavax and Janssen mRNA sequences with artificially added substitutions R682S/Q, R683Q, R685G/Q, K986P, and V987P.

**Figure 4.**
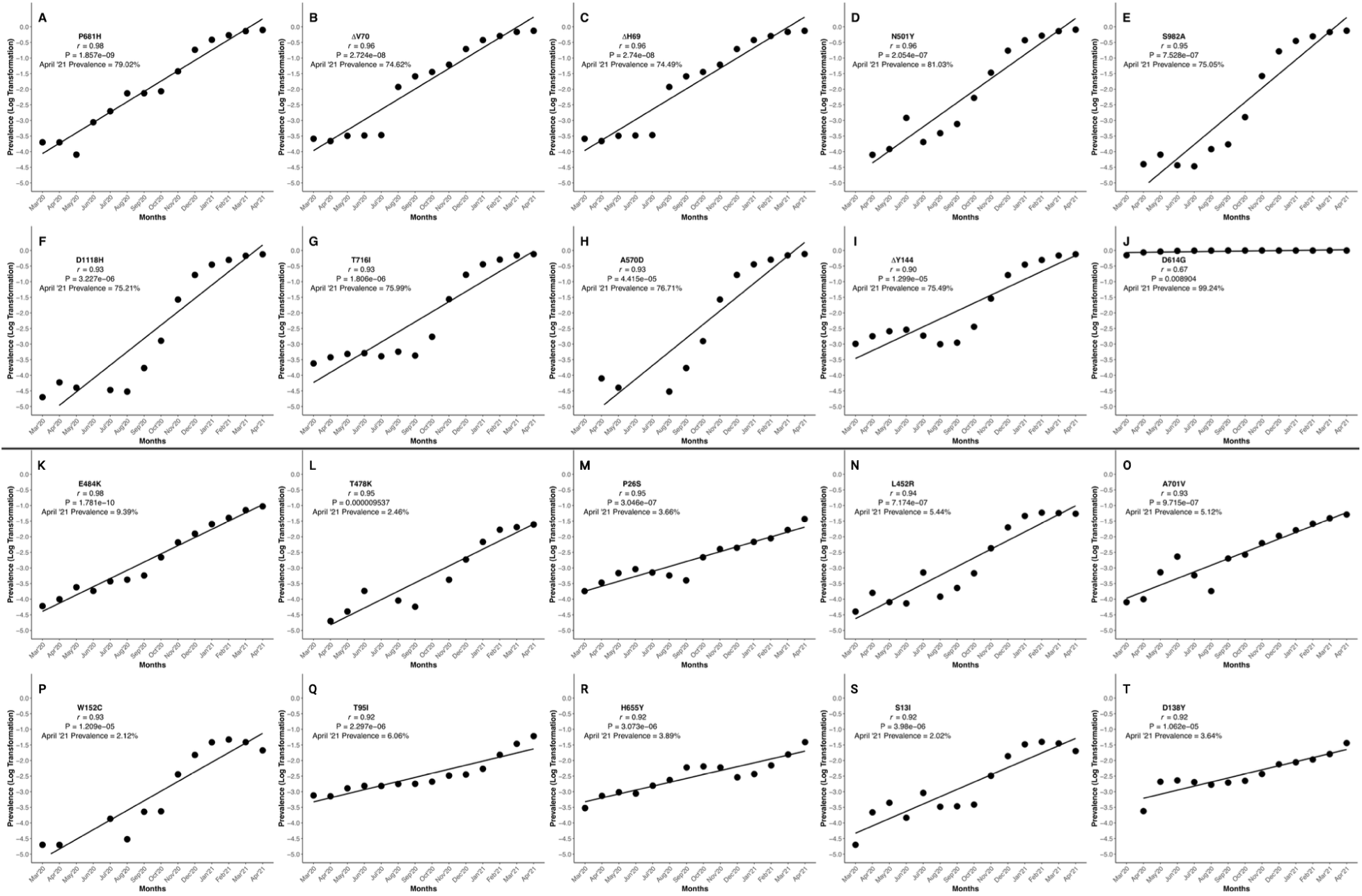
Pearson’s Correlation of Logarithmically-Transformed Prevalence Ratios of the Most Emergent SARS-CoV-2 Mutations of Concern and Interest Selected via the Algorithm. This figure shows the graphical representation of the logarithmically-transformed prevalence data used to calculate the Pearson’s correlation of each of the twenty most emerged (of concern) and emergent (of interest) spike protein substitutions and deletions. The substitutions and deletions of concern here are in order of decreasing *r* value, and each has a unique alphabet identifier A) P681H, B) ΔV70, C) ΔH69, D) N501Y, E) S982A, F) D1118H, G) T716, H) A570D, I) ΔY144, and J) D614G. The algorithm uses the monthly prevalence data from these ten spike protein substitutions and deletions, and they are the most concerning of all spike changes. The substitutions and deletions of interest here are in order of decreasing *r* value and each unique substitution or deletion is denoted by a letter of the English alphabet, K) E484K, L) T478K, M) P26S, N) L452R, O) A701V, P) W152C, Q) T95I, R) H655Y, S) S13I, and T) D138Y. Graphs were generated using RStudio version 1.3.1093 (R version 4.0.3) and the ggplot2 package. Graphs were compiled and the final figure generated using Biorender.com.

**Table 3.**
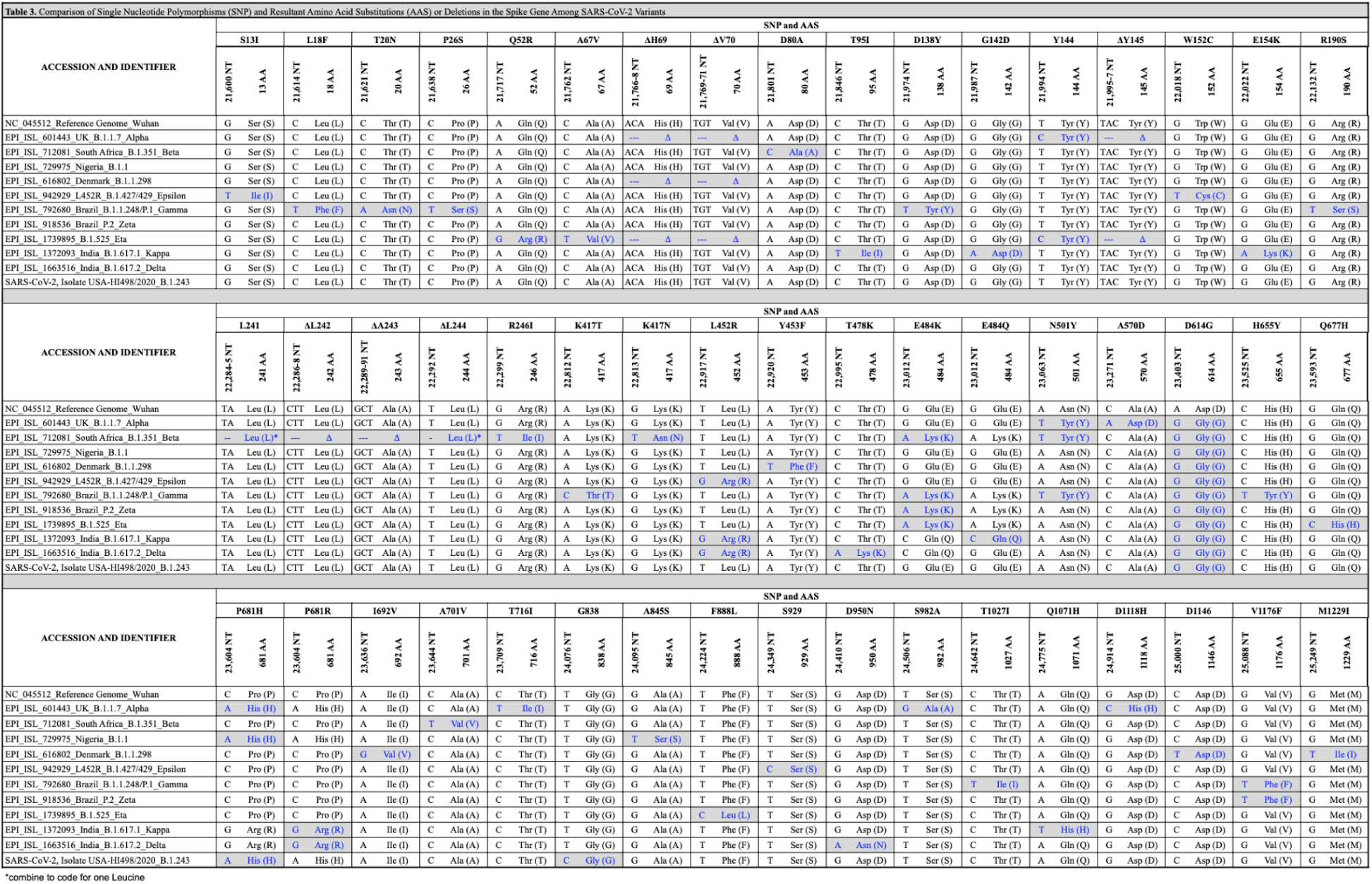
Comparison of Single Nucleotide Polymorphisms (SNP) and Resultant Amino Acid Substitutions (AAS) Among SARS-CoV-2 Variants

The PubMed search for epitope predictions returned 42 publications. In total, 393 *in silico* predicted B and T cell epitopes corresponding to the spike protein were mapped from these publications (Figure 3B). Of these, 108 epitopes involved the N-terminal domain (NTD), 102 epitopes involved the receptor binding domain (RBD), 7 epitopes involved the S1/S2 furin cleavage site, 10 epitopes involved the fusion peptide, 20 involved heptad repeat 1, 12 involved heptad repeat 2, 12 involved the transmembrane region, and 8 involved the intracellular tail domain. The remaining 112 epitopes fell outside of these domains. Further, 239 and 151 epitopes were in the S1 and S2, respectively, with at least one predicted epitope covering 97% of the spike protein.

### Variant Comparison

The unique nucleotide mutations and resulting AAS and deletions for each of the twelve SARS-CoV-2 variants, in comparison to the reference sequence, are shown in Table 3. BNT162b2 (Pfizer) and mRNA-1273 (Moderna) vaccines both contain two AAS (K986P and V987P) (Figure 3C.xii).^52^ Novavax and Janssen vaccines also contain two AAS, K986P and V987P. Further, additional AAS at the furin cleavage site includes, R862Q, R683Q, and R685Q (Novavax), and R682S and R685G (Janssen) (Figure 3C.xiii).^52^

### B.1.243 Phylogeny and Origin Tracking

The Hawai’i SARS-CoV-2 sequences in the GenBank and GISAID were combined with all worldwide B.1.243 lineages to produce an initial MAFFT alignment of 8,820 sequences. Further, 4,273 sequences with ambiguities and 1,596 duplicate sequences were removed. The final alignment for constructing the phylogenetic tree was 2,953 unique and unambiguous B.1.243 sequences. Using this method,^13^ we were able to define the origin of SARS-CoV-2, Isolate USA-HI498 2020 and HI708 (Figure 5).

**Figure 5.**
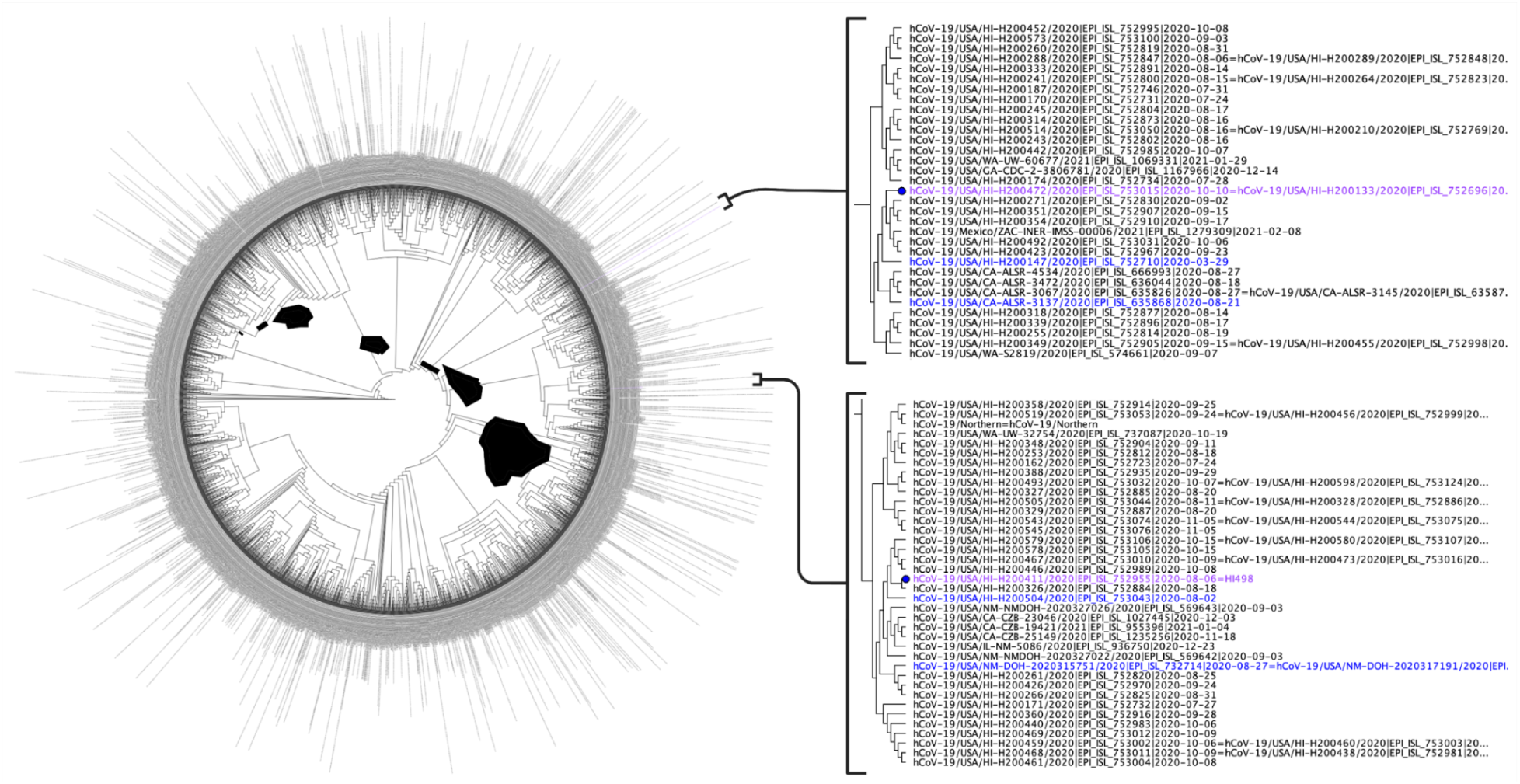
Phylogenetic Tree of all B.1.243 Lineage Sequences Worldwide. This figure displays the phylogenetic tree used to determine the origin of the B.1.243 sequences used in this study. We use 8,822 SARS-CoV-2 B.1.243 whole-genome sequences published in the Global Initiative on Sharing Avian Influenza Data (GISAID) and GenBank as of April 12, 2021 to define the origin. From the 8,822, 4,273 had ambiguous nucleotides between the 5’ and 3’ untranslated regions as determined using multiple alignment using fast Fourier transform (MAFFT). Further, 1,588 were duplicate sequences and eight had duplicate identifications as determined by the sRNA Toolbox. Therefore, the final tree was constructed using FastTree in Geneious Prime 2021.1.1 (http://www.geneious.com) from 2,953 unique and unambiguous SARS-CoV-2 whole-genome sequences. The HI498 (purple text) origin is defined as New Mexico (blue text) and the HI-708 (purple text) origin is defined as California (blue text).

## Discussion

In this report, we lay the foundation for an adaptive and rational algorithm for monitoring SARS-CoV-2 evolution, quantitating variants, substitutions, and deletions, in the context of the vaccine design. Further, we describe the isolation, genetic characterization, phylogenetic analysis, and immunogenetic epitopes of the spike protein based on the SARS-CoV-2 lineage B.1.243 from Hawai’i. We employed B.1.243 to establish and validate the algorithm, as well as analyze VOC and VOI in the context of emerging spike protein amino acid changes for surveillance and future vaccine design.

### Hawai’i Isolate B.1.243

Hawai’i has not been spared from this pandemic, and the Pacific Islander population here is disproportionately infected with SARS-CoV-2 as compared to other races and ethnicities.^12^ Following isolation and identification of the B.1.243 lineage from the isolate SARS-CoV-2, Isolate USA-HI498 2020, and virus strain HI708, we found through curation and analysis of published sequences that the B.1.243 lineage was once the dominating lineage in Hawai’i, causing more than 40% of all cases. The Hawai’i B.1.243 lineage is not likely to escalate into a VOC, as the prevalence has decreased worldwide over the past several months. However, the disproportionate infection rates among the Pacific Islander population remains.^12^

The B.1.243 lineage that once dominated in overall prevalence in Hawai’i was introduced from Washington, California, Pennsylvania, and New Mexico, with the majority of sequences arising from horizontal transfer within Hawai’i. SARS-CoV-2, Isolate USA-HI498 2020 was introduced in Hawai’i from New Mexico and the HI-708 SARS-CoV-2 strain originated from California. As this report is written, similar to the continental United States the SARS-CoV-2 Delta variant is rapidly spreading in Hawaii.^53^

### Quantitation and Analysis of Variants of Concern

At the time of the submission, the CDC identifies four SARS-CoV-2 variants as VOC: B.1.1.7 (Alpha), B.1.351 (Beta), B.1.617.2 (Delta), and P.1 (Gamma).^1^ Our quantitative data analysis supports the exponential emergence of these VOC, with the most emergent being P.1, followed by B.1.617.2, B.1.351, and B.1.1.7, with Pearson’s correlation *r*-values of 0.97, 0.96, 0.94, and 0.92, respectively. The quantitative analysis described in this report gives a numerical value to each VOC emergence, predicting the likelihood that the lineage will become prevalent and spread through the population. This value then indicates which VOC genomes are likely to possess evolutionarily selective changes. Previously, using this quantitative analysis, we have demonstrated that the P681H substitution, which had a prevalence of 2% worldwide in December 2020, then emerged to 79% in April 2021.^14^ Similarly, using this quantitative analysis in April 2021, we predicted the spread of the Delta variant in Hawaii and worldwide as of June 2021.^13^

In this report we demonstrate the characteristic emergence and selective evolution of VOC using the B.1.1.7 VOC as an example, with a Pearson’s correlation of 0.92. The B.1.1.7 VOC, the most prevalent VOC worldwide in April 2021, has spread across the globe after emerging in the United Kingdom in December 2020.^31^ As stated, the B.1.1.7 VOC represents the prototypic VOC which has evolved continuously by evading vaccine sera and becoming more transmissible.^54, 55^ Similarly, the Delta variant, predicted to be exponentially emerging by this quantitative analysis with an *r-*value of 0.96 as of April 2021 at 1% prevalence, has become the most prevalent worldwide as of June 2021, representing 64% of worldwide sequences.

The virus transmissibility, prevalence, and decrease of treatment and vaccine efficacy is concerning. Each of these viral properties has been attributed to specific amino acid alterations in the spike protein. For example, the L452R substitution, prevalent in many VOC and VOI, causes a two-fold increase in viral shedding and renders multiple FDA approved monoclonal antibody treatments ineffective.^56^ Thus, the quantitation of VOC leads to identification and quantitation of their respective mutations. The pairwise heat map between variants and mutations (Table 2) indicates that variants do not necessarily evolve with mutations, and the genomes may spontaneously acquire or revert to wildtype, as demonstrated in other statistical analysis for monitoring this pandemic.^57^

### The Algorithm

We were able to establish the algorithm described in this report based on the ten most emerged AAS and deletions. We evaluated nine of the AAS and deletions observed among the variants in this study, as of May 12, 2021, with *r* > 0.9 and >70% prevalence in April 2021. These AAS and deletions included, P681H, ΔV70, ΔH69, N501Y, S982A, D1118H, T716I, A570D, and ΔY144. The data show that the average time for an emerging substitution (*r* > 0.9) to go from >30% monthly prevalence percentage to >50% prevalence is 2.25 months. Extrapolating these findings, the timeframe for Pfizer to identify emerging changes and manufacture them into a new version of the BNT162b2 vaccine would be 60-110 days. This timeframe is roughly equivalent to the algorithm’s predictive value.^58^ Additionally, the tenth AAS used in the development of the algorithm was D614G that allowed us to discern the previous month prevalence, which is an important parameter, as once an emerging mutation is established in the genome the *r* value will decrease considerably. The nine substitutions and deletions also display an average of 4.75 months from >2% monthly prevalence to >50% monthly prevalence. Therefore, *r* > 0.9 and a prevalence of >2% is sufficient to establish a mutation as being of interest, whereas 30% prevalence escalates a mutation to the status of concern. From the evaluation of the ten total spike protein changes, the algorithm concludes that an *r* > 0.9 and a >30% prevalence percentage is an optimal time to classify a substitution as concerning, and consider the substitution for inclusion into vaccine primary structure for 60 day production time.

These mutations of interest and concern can also serve to facilitate and focus research using infectious clone^59^ and pseudoviruses^60^ to determine the functional characteristics. Amino acid substitutions are responsible for, i) changing epitopes at a level to evade antibodies,^1, 61, 62^ ii) allowing the virus to localize anatomically,^63^ iii) increasing viral shedding,^62^ iv) increasing binding affinity,^64^ v) giving the virus binding protein the ability to change conformations more efficiently,^65^ and vi) causing diagnostic false negatives.^66^ Therefore, of great importance in vaccine development is identifying which spike protein changes are most prevalent worldwide across all sequenced genomes. Each substitution or deletion, or combination thereof, could potentially serve as an epitope as shown in the epitope map (Figure 3C). Booster vaccines using this algorithm can therefore prepare the vaccinated for any variant they are most likely to encounter by identifying and including the emerging and emerged amino acid changes representing the majority of SARS-CoV-2 worldwide.

### In silico Predicted Immunogenic Epitopes in Relation to Variants of Concern

Epitopes are found across nearly the entire spike protein. The *in silico* compilation of predicted B cell and T cell epitopes demonstrated that 53% (210/393) of all epitopes occur in the NTD and RBD. This is consistent with an *in vivo* mRNA-LNP vaccine study that found a vast majority of the CD8+ T cell response target epitopes in the N-terminal portion of the Spike protein.^67^ The same study found that CD4+ T cell responses target both S1 and S2.^67^ Additionally, the majority of variant mutations and neutralizing antibody targets occur in S1.^55^ As S1 is shed in the coronavirus model of fusion,^68^ and S2 is responsible for fusion, the diversity of S1 in SARS-CoV-2 and the epitope targeting concentration in S1, indicate that SARS-CoV-2 vaccines will need to adapt along with the virus.

### Current Vaccines, Structures, and Introduced Mutations in Relation to Variants of Concern

The Pfizer/BioNTech and Moderna/National Institute for Allergy and Infectious Diseases (NIAID) are mRNA vaccines that both use the S gene of SARS-CoV-2 S-2P. S-2P is the reference sequence with the substitution of two prolines (K986P, V987P) to stabilize the pre-fusion conformation of the spike protein.^52^ The Janssen and Novavax vaccines for SARS-CoV-2 are Ad26-vectored and protein-based, respectively, and use the S-2P sequence with the addition of three AAS (R682S/Q, R683Q, R685G/Q) at the furin cleavage site.^52^ These furin AAS have been shown to further stabilize the pre-fusion spike conformation, and increase neutralizing antibody titers.^52^ The majority of other vaccines, including whole virus vaccines, S protein vaccines, and RBD vaccines do not utilize the S-2P method of pre-fusion stabilization.^52, 68^ Current post-vaccination sera has reduced neutralization against the VOC as the COVID-19 pandemic proceeds into endemicity. As the loss of neutralization is observed against the most emerging VOC and VOI herein, there is a need for a new generation of vaccines for optimal efficacy in years to come.

### Future Directions

Our findings have relevance to the future of tracking SARS-CoV-2 and of SARS-CoV-2 vaccine design. The future SARS-CoV-2 vaccines are akin to influenza virus vaccines. That seeming nature is that influenza vaccines change yearly depending on the previous years surveillance data.^69^ If established in real-time, the herein described algorithm will allow researchers to understand the evolving SARS-CoV-2 genome preemptively rather than responsively.

As the number of worldwide SARS-CoV-2 sequences swells into many millions between GenBank and GISAID, the need for an Application Programming Interface (API) between the two databases is needed now. Such an API would allow the quantitation of emerging mutations and variants to alarm when necessary, rather than at arbitrary media discretion. Real-time quantitation would then allow vaccines to evolve preemptively. As things stand, the data is self-diversifying by researchers’ choice of submission to one database or the other, rendering GISAIDs mutation filters less informative on a worldwide scale. As one database headquarters in Germany (GISAID) and the other in the United States (GenBank), the world needs solidarity now more than ever as we combat this global pandemic.

## Conclusions

Here, we isolate SARS-CoV-2 in Hawai’i and evaluate the WGS of these SARS-CoV-2 isolates. We apply our archetype method for predicting exponentially emerging mutations to these isolates, then evolve the quantitative analysis into an algorithm to evaluate the VOC, VOI, and their mutations, allowing further classification of mutations as concern and interest. This algorithm can now serve as a baseline for choosing the primary structure of vaccines. Additionally, we graphically compare the S gene of the Hawai’i isolate with SARS-CoV-2 VOCs, predict the emergence of SARS-CoV-2 S gene mutations and protein substitutions and deletions, and evaluate these substitutions and deletions in the context of epitopes. In conclusion, we create a foundation for future SARS-CoV-2 monitoring and vaccine efforts as we move forward in this pandemic (Figure 6), and demonstrate the need for sequence database solidarity. These pandemic efforts cannot remain in the context of being responsive and reactive, but need also to be preemptive and predictive. Preemptive and predictive is possible with the approach herein.

**Figure 6.**
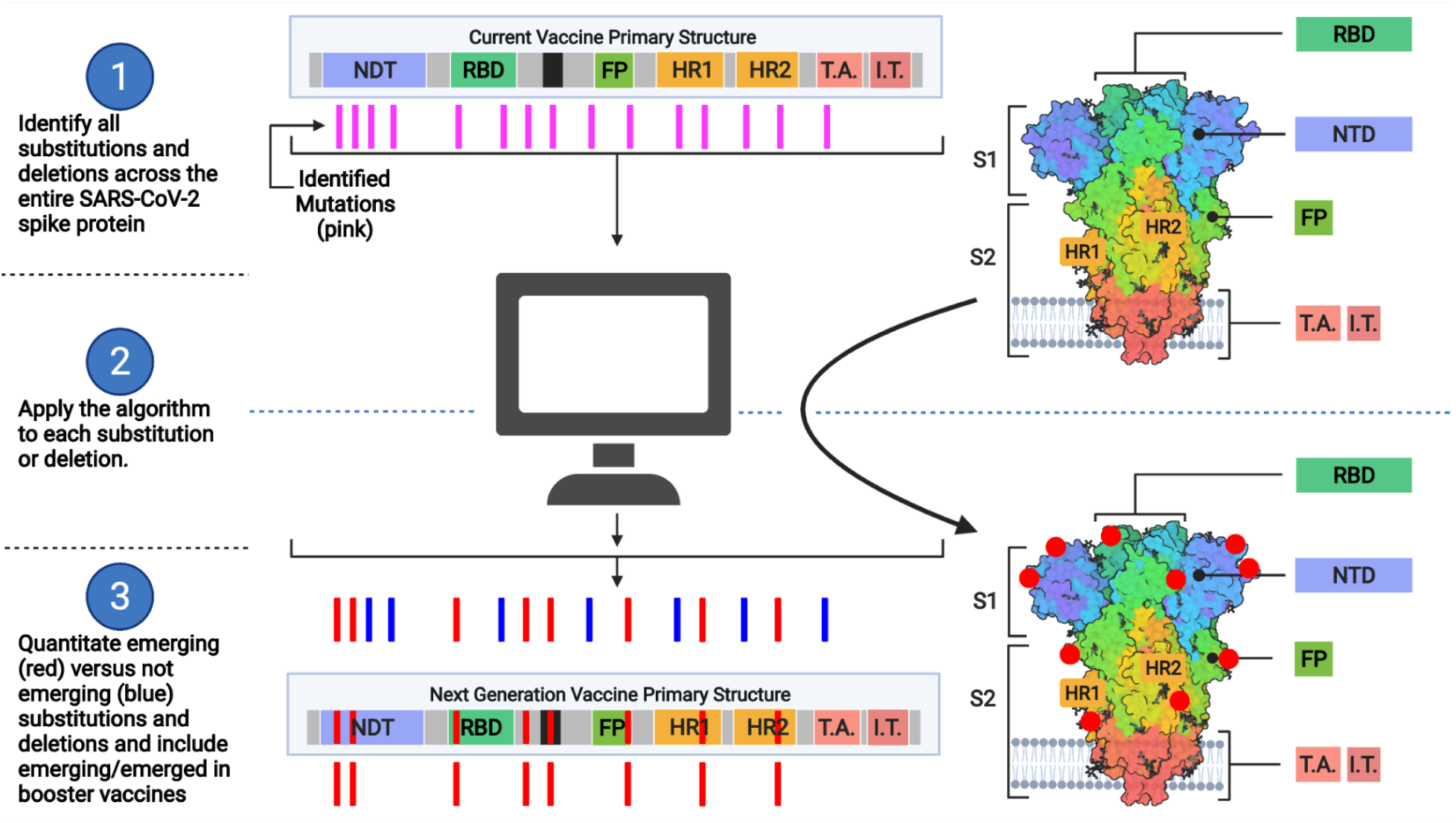
Applying the Algorithm to Vaccine Design. The model displays how the algorithm would lead to the design of next-generation vaccines based on the S gene. Part one of the model identifies all SARS-CoV-2 genomic mutations and spike amino acid substitutions and deletions (represented as pink lines) based on the worldwide sequence databases GISAID and GenBank. The current vaccine design as it translates into proteins is depicted in the top center and top right. Part two of the model will apply the quantitative analysis and algorithm described in this report to each of the protein changes identified in part one. The quantitative analysis determines emergence via logarithmic transformation of prevalence and Pearson’s correlation. The algorithm then applies criteria to the quantitative analysis and previous months prevalence for determining which changes are likely to be in the majority of SARS-CoV-2 for incorporation in the next-generation vaccine. Part three of the model determines which substitutions and deletions are exponentially emerging or emerged (red lines) and which are not (blue lines). From part three, the mRNA sequence of vaccines can then incorporate the emerging and emerged mutations so that the folded protein (bottom right) will contain the protein changes (red dots) most prevalent worldwide by the time the next-generation vaccine is manufactured and administered. As a result, these substitutions and deletions will present the most appropriate epitopes of the SARS-CoV-2 spike protein to vaccine recipients.

## Acknowledgements

This research was supported by a grant (P30GM114737) from the Pacific Center for Emerging Infectious Diseases Research, COBRE, a grant (P20GM103466-20S1) from the INBRE, National Institute of General Medical Sciences, and a grant (U54MD007601) from Ola Hawaii, National Institute on Minority Health and Health Disparities, NIH. The H051 clinical trial is registered at ClinicalTrials.gov (#NCT04360551). Computation was supported by NSF grant #1920304 on the University of Hawai’i MANA High Performance Computing Cluster. The viral genome sequences used in this publication are publicly available from GenBank (https://www.ncbi.nlm.nih.gov/sars-cov-2/) and GISAID (https://gisaid.org). Tables of acknowledgements for the genome sequences from GISAID are available at: https://github.com/dpmaison/Algorithm-for-the-Quantitation-of-Variants-of-Concern-for-Rationally-Designed-Vaccines. Other genome sequences from GISAID are referenced in-text. We thank Dr. Jennifer Saito at the Advanced Studies in Genomics, Proteomics and Bioinformatics (ASGPB) Core, UHM for assistance with WGS.

---

**Supplementary Figure 1.**
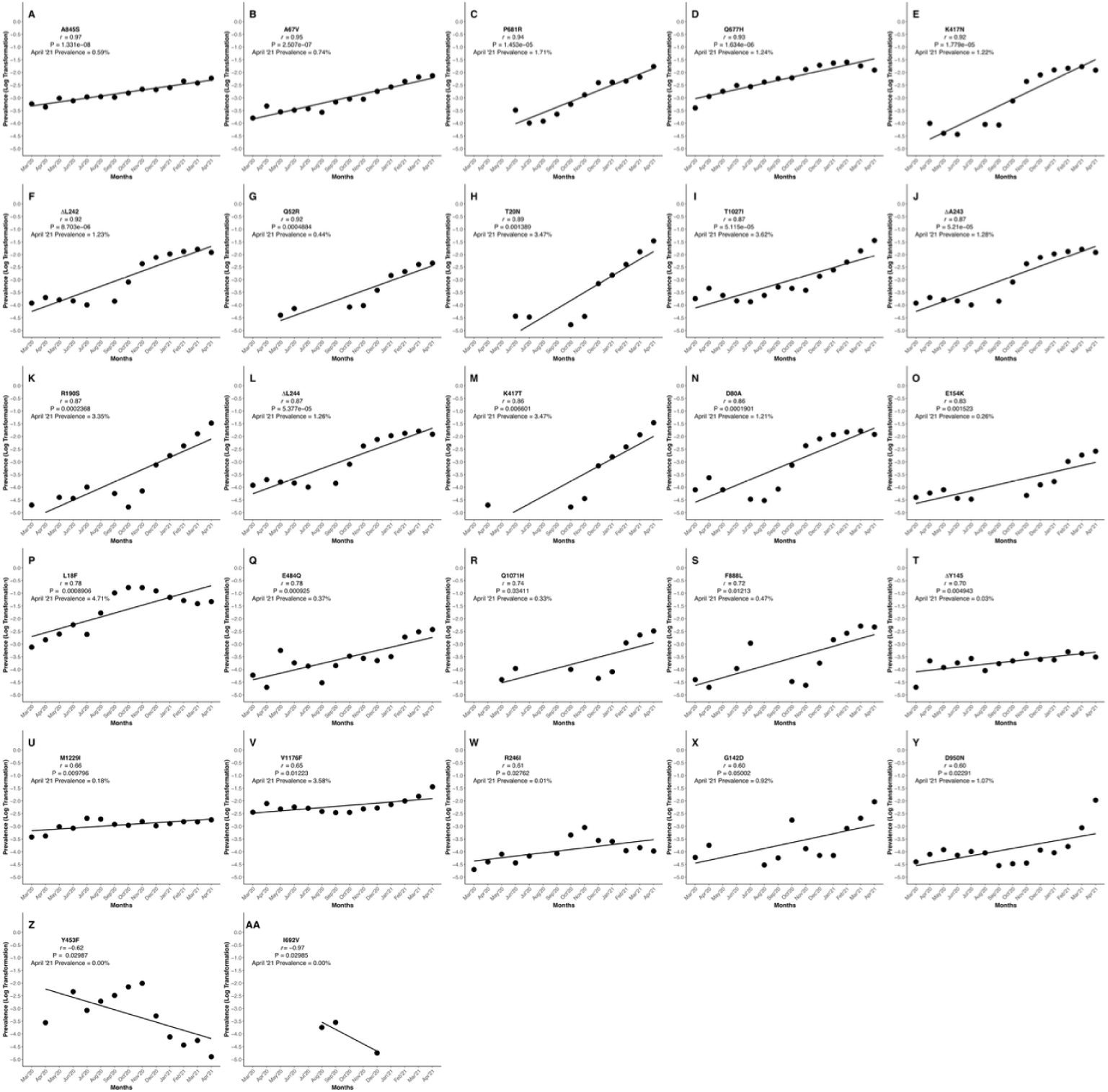
Pearson’s Correlation on Logarithmically-Transformed Prevalence Ratios of the Remaining SARS-CoV-2 Variant Amino Acid Substitutions and Deletions Not Currently Selected via the Algorithm. This figure shows the graphical representation of the SARS-CoV-2 spike protein amino acid substitutions and deletions not currently concerning due to low previous month prevalence, low *r* value, or insignificant P value. Though not yet of concern as of April ‘21, those substitution and deletions represented by high *r* values should be cause for close monitoring. Each graph denoted by an alphabetical character or characters represents a unique amino acid substitution or deletion in the spike protein of SARS-CoV-2 (A) A845S, B) A67V, C) P681R, D) Q677H, E) K417N, F) ΔL242, G) Q52R, H) T20N, I) T1027I, J) ΔA243, K) R190S, L) ΔL244, M) K417T, N) D80A, O) E154K, P) L18F, Q) E484Q, R) Q1071H, S) F888L, T) ΔY145, U) M1229I, V) V1176F, W) R246I, X) G142D, Y) D950N, Z) Y453F, and AA) I692V). Graphs were generated using RStudio version 1.3.1093 (R version 4.0.3) and the ggplot2 package. Graphs were compiled and the final figure generated using Biorender.com.

## Notes

### Competing Interest Statement

The authors have declared no competing interest.

