## Supplementary figures and images for "Algorithm for the Quantitation of Variants of Concern for Rationally Designed Vaccines Based on the Isolation of SARS-CoV-2 Hawaiʻi Lineage B.1.243"

### Supplementary Figure 1

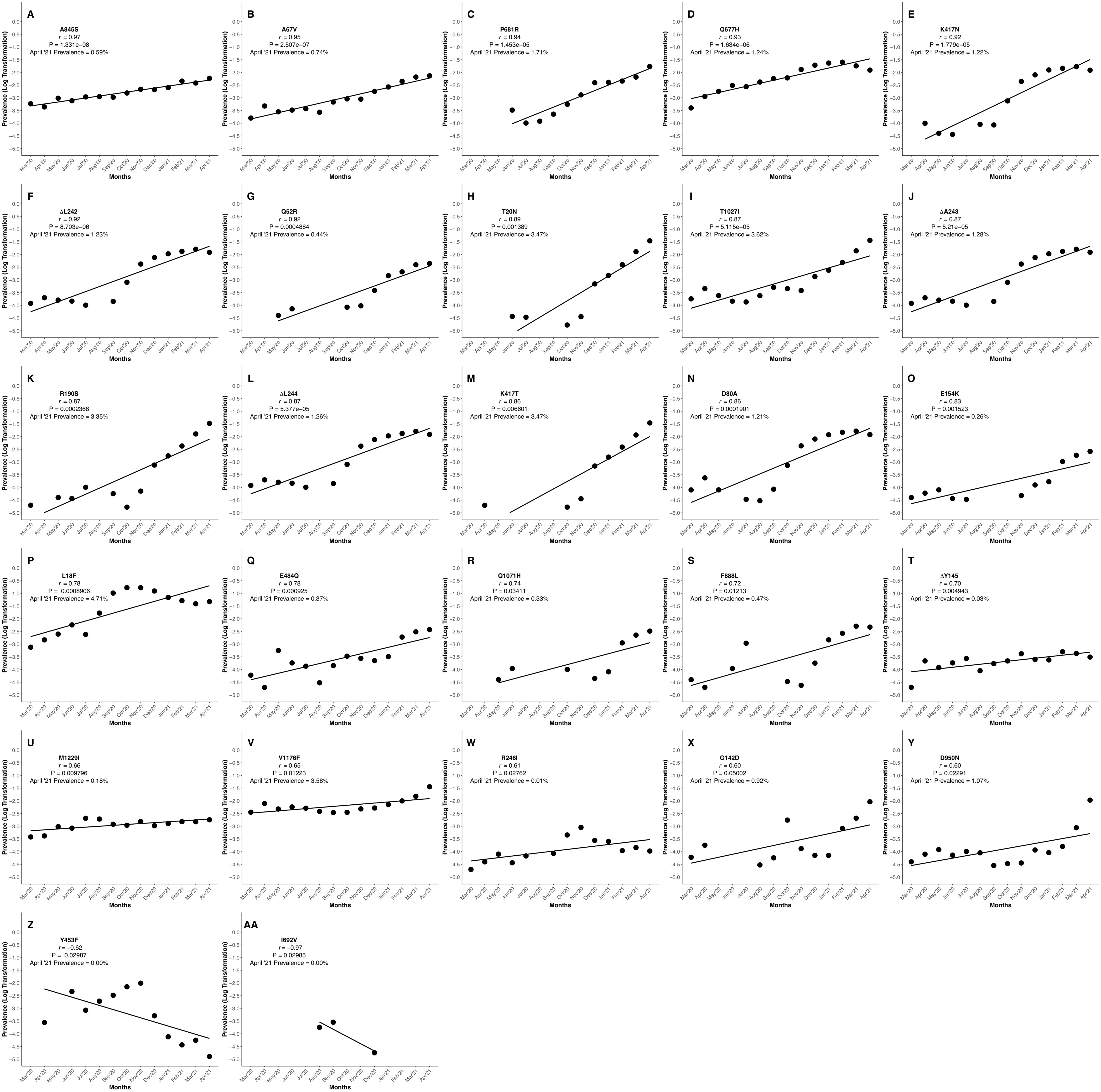
